# Intestinal stem cell renewal controlled by capillary morphogenesis gene 2 following injury

**DOI:** 10.1101/2025.01.07.631493

**Authors:** Lucie Bracq, Audrey Chuat, Béatrice Kunz, Olivier Burri, Romain Guiet, F. Gisou van der Goot

**Affiliations:** Global Health Institute, School of Life Sciences, EPFL, Lausanne Switzerland; BioImaging and Optics Core Facility, School of Life Science, EPFL, Lausanne Switzerland

## Abstract

Patients with the rare genetic disorder Hyaline Fibromatosis Syndrome (HFS) often succumb before 18 months of age due to severe diarrhea and protein-losing enteropathy. As HFS is caused by loss-of-function mutations in the gene encoding capillary morphogenesis gene 2 (CMG2), also known as Anthrax Toxin Receptor 2, these symptoms highlight a critical yet unclear role for CMG2 in the gut. Here, we demonstrate that CMG2 knockout mice exhibit normal colon morphology and no signs of inflammation until the chemical induction of colitis. In these conditions, the colons of knockout mice do not regenerate despite previously experiencing similarly severe colitis, due to an inability to replenish their intestinal stem cell pool. Specifically, CMG2 knockout impairs the transition from fetal-like to Lgr5+ adult stem cells, which is associated with a defect in ß-catenin nuclear translocation. Our findings suggest that CMG2 functions as a context-specific modulator of Wnt signaling, essential for replenishing the pool of intestinal stem cells following injury. This study provides new insights into the molecular mechanisms underlying protein-losing enteropathy in HFS and offers a broader understanding of fetal-like regenerative responses.

**GRAPHICAL ABSTRACT:** 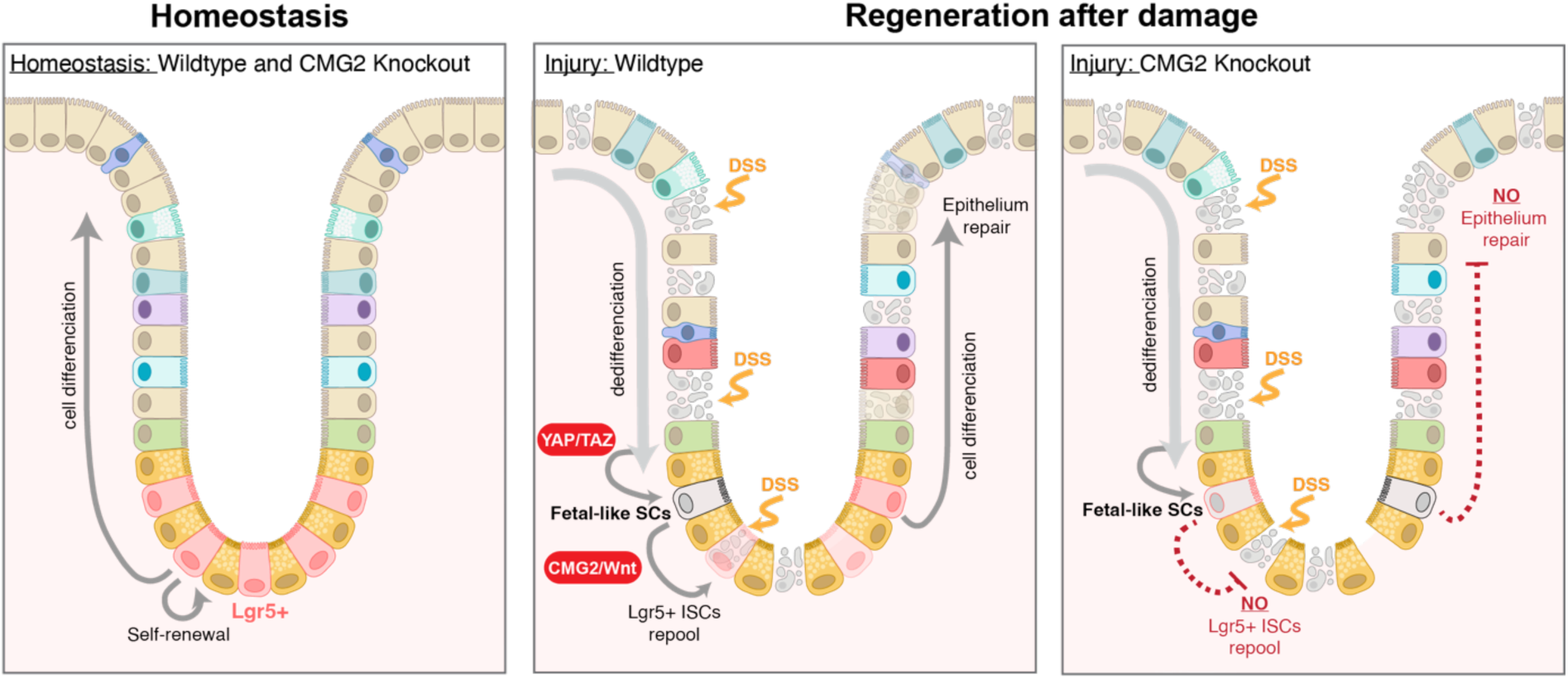

## INTRODUCTION

Hyaline Fibromatosis Syndrome (HFS; OMIM #228600) is a rare autosomal disorder that, in its most severe form, leads to death before 18 months of age due to intractable diarrhea or recurrent pulmonary infections^1^. Despite the critical nature of these symptoms, there are currently no effective interventions, primarily because the underlying molecular mechanisms remain poorly understood. The defining feature of HFS is however the abnormal accumulation of hyaline material, particularly collagen VI^2^, in the skin, which results in large subcutaneous nodules, gingival hypertrophy, and painful joint contractures^3–9^. HFS is caused by mutations in the Capillary Morphogenesis Gene 2 (CMG2), with evidence showing that CMG2 regulates extracellular collagen VI levels^2^. However, the severe diarrheal symptoms and associated intestinal lymphangiectasia, often culminating in fatal protein-losing enteropathy¹, suggest that CMG2 may play a distinct and crucial role in intestinal physiology that has not yet been characterized. This study aims to address this critical gap by elucidating the role of CMG2 in the gut and uncovering the mechanisms responsible for the devastating intestinal symptoms observed in severe HFS patients.

CMG2, also known as ANTXR2, controls the turnover of collagen VI by mediating its intracellular uptake and degradation in lysosomes^2^. CMG2, a type I membrane protein featuring an extracellular von Willebrand A (vWA) domain, can indeed bind extracellular matrix components, including laminin and types IV and VI collagen ^10^, as well the actin cytoskeleton and actin modulators intracellularly^11^. Upon CMG2 loss of function, multi systemic-deposition of collagens, in particular Collagen VI, occurs in HFS patients^2,11–13^. CMG2 may have roles other than controlling the abundance of the ECM. In particular, while the intestine of two infants with severe HFS showed accumulation of ECM and in particular collagen VI, no primary defect in the function of the intestinal epithelium was observed based on in vitro culture of patient cells either as gut organoids or intestinal epithelial layers^13^.

We therefore speculate that CMG2 does not act directly on intestinal epithelial cells but plays a role in crucial processes such as homeostatic cellular turnover and intestinal regeneration. The YAP and Wnt signaling pathways are central to these processes^14–16^, and several studies have suggested links between CMG2 and these pathways^2,17–21^. The normal rapid cellular turnover of the intestinal epithelium is controlled by the interplay between Wnt, Notch, and YAP/Taz signaling pathways^22^. After intestinal injury, intestinal stem cells (ISCs) are depleted^23–29^, triggering a regenerative response to restore tissue integrity^14,22,30–32^. Recent research indicates that this regeneration involves dedifferentiation and a fetal-like regenerative response^14,23,25,26,31–42^, a process highly dependent on YAP and Wnt signaling interactions^25,43,44^. However, the precise mechanisms governing this regenerative response remain incompletely understood.

In this study, we utilized a knockout mouse model to investigate the role of CMG2 in the colon under both normal and regenerative conditions. While no abnormalities were observed in the intestinal architecture or function under homeostatic conditions, the scenario changed following chemically-induced colitis^45,46^. *Cmg2* knockout (*Cmg2*^KO^) mice failed to restore intestinal integrity, underscoring a critical role for CMG2 in regeneration. Our analysis showed that CMG2 is not involved in the initial fetal-like reversion phase, but is essential to replenish the Lgr5+ intestinal stem cell (ISC) pool— an important Wnt-dependent step for epithelial proliferation and barrier restoration. These findings highlight CMG2’s vital role in Wnt-driven ISC replenishment during intestinal regeneration, providing new insights into the transition from fetal-like to adult ISC states, and provide molecular understanding for the chronic diarrhea seen in severe HFS patients.

## RESULTS

### Gut in *Cmg2*^KO^ mice is normal under basal conditions but fails to recover from DSS-induced colitis

To explore the role of CMG2 in intestinal homeostasis, we used our previously described *Cmg2*^KO^ mice^2^. Using this model, we found no obvious symptoms associated with intestinal dysfunction (Fig. S1a), nor abnormalities in crypt architecture detectable by histological examination (Fig. S1b-d) in *Cmg2*^KO^ mice. We then specifically measured collagen VI, as it has been observed to accumulate over time in both HFS patient nodules and *Cmg2*^KO^ mouse tissues^2^, due to impaired CMG2-mediated turnover. However, the colon of 8-week-old *Cmg2*^KO^ mice had normal levels of collagen VI as measured by Western blot analysis of protein extracts (Fig. S1e-f), histological examination (Fig. S1b), immunofluorescence (Fig. S1g-h), and qPCR (Fig. S1i). Additionally, the loss of CMG2 did not affect the number of ISCs and actively proliferating cells, both essential for maintaining epithelial integrity under homeostatic conditions^47–49^. Markers for ISCs, including Lgr5 mRNA (Fig. S1j-l), and indicators of cell proliferation (Fig. S1m-o) also showed no significant differences between *Cmg2*^KO^ and *Cmg2*^WT^ mice, suggesting normal cellular turnover. Furthermore, *Cmg2*^KO^ mice exhibited no signs of inflammation (Fig. S1p). Overall, these findings indicate that the loss of CMG2 does not significantly impact the architecture or function of the mouse colon under homeostatic conditions.

To explore this seeming discrepancy between the healthy colons in *Cmg2*^KO^ mice versus the recurrent, non-infectious diarrhea seen in HFS patients, we examined whether differences between control and KO mice would emerge under stress conditions. Using the well-established dextran sodium sulfate (DSS)-induced colitis model (Fig. 1a-b), both *Cmg2*^KO^ mice and their wild-type littermates received 3% DSS in their drinking water for 7 days, consuming similar amounts (Fig. 1c). In both groups, the treatment led to established effects of DSS-induced colitis^45,46,50,51^, including body weight loss (Fig. 1d), diarrhea, and rectal bleeding (Fig. 1e). To evaluate potential differences in colitis severity, we monitored the disease activity index (DAI)^45,46,50^ daily throughout the experiment (Fig. 1d-f). Until the end of the 7-day DSS treatment, both groups showed a similar, progressive increase in DAI (Fig. 1f) alongside a sharp rise in the intestinal inflammation marker lipocalin-2 (Fig. 1g), confirming the successful induction of colitis in our model. Histological examination immediately after the DSS challenge revealed significant colon alterations in both *Cmg2*^WT^ and *Cmg2*^KO^ mice, including loss of crypt architecture (Fig. S2a), depletion of epithelial cells (Fig. S2a,b), and reduced cell proliferation (Fig. S2a,c). DSS treatment also led to the loss of Lgr5^+^ stem cells (Fig. S2a,d-e), a common feature of intestinal damage, with similar levels of inflammation observed in both groups (Fig. S2f-i). Overall, these results indicate that CMG2 does not affect the severity of DSS-induced colitis.

**Figure 1:**
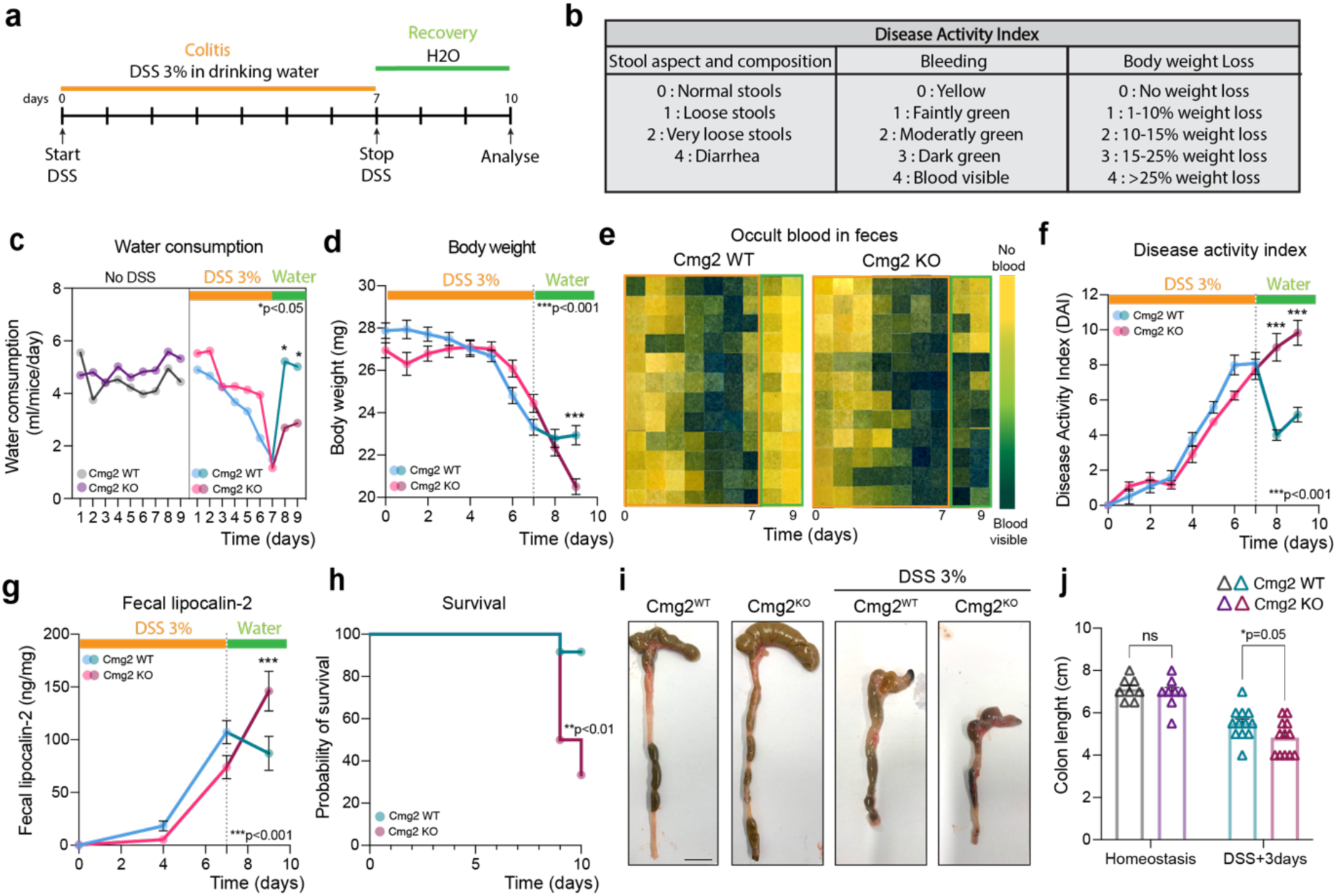
*Cmg2*^KO^ mice fail to recover from DSS-induced colitis. 8-week-old *Cmg2*^WT^ or *Cmg2*^KO^ males were given 3% Dextran-Sulfate-Sodium (DSS) in their drinking water for 7 days, then switched to regular drinking water for 3 days to recover. **(a)** Experimental scheme of DSS-induced colitis. **(b)** Disease activity index (DAI) scoring and **(c)** water consumption was measured daily during the 10-day experiment. In (c), results are mean ± SEM. Each dot represents the mean of n = 2 or 3 different cages. *P* values were obtained using a two-way ANOVA Tukey’s multiple comparisons test. **(d)** Body weight loss, **(f)** the aspect of the feces and presence of occult blood were monitored and used for the **(e)** DAI. Results are mean ± SEM. Each dot represents the mean of n = 12 mice per genotype. *P* values were obtained by two-way ANOVA Šídák’s multiple comparisons test. In (e), presence of blood in feces was detected using a urine strip test from collected feces. Each row corresponds to a single mouse and each column to a different timepoint. Feces were collected daily, and **(g)** fecal lipocalin-2 was quantified by ELISA. Results are mean ± SEM. Each dot represents the mean of n = 12 mice per genotype. *P* values were obtained using a two-way ANOVA Šídák’s multiple comparisons test. Excessive body weight loss (>25%) required animal euthanasia as shown in **(h)** probability of survival. On day10, mice were euthanatized, organ collected and **(i-j)** colon length were measured. Representative images of n = 8 mice per genotype for untreated mice and n = 12 mice per genotype for DSS-treated mice. Scale bar, 1 cm. In (J), Results are mean ± SEM. Each symbol represents a single measurement. *P* values were obtained using an unpaired T-test.

After the 7-day DSS treatment, mice were switched to normal drinking water for 3 days, a period known to allow intestinal regeneration in control mice^45,46,50^ (Fig. 1a). During this phase, *Cmg2*^WT^ mice showed typical signs of recovery, including a stabilization of body weight (Fig. 1d), cessation of rectal bleeding (Fig. 1e), reduction in DAI (Fig. 1f), and reduced levels of lipocalin-2 in feces (Fig. 1g). In contrast, *Cmg2*^KO^ mice continued to lose weight (Fig. 1d), ultimately reaching the legal euthanasia limit (>25% body weight loss), which led to a sudden increase in apparent mortality (Fig. 1h). Moreover, both occult blood and elevated lipocalin-2 levels persisted in the feces of *Cmg2*^KO^ mice (Fig. 1e). Additionally, colons from *Cmg2*^KO^ mice euthanized on day 10 were significantly shorter than those of their wild-type counterparts (Fig. 1i-j). Together, these findings indicate that while CMG2 does not affect the initial severity of DSS-induced colitis, it plays a crucial role in promoting intestinal recovery after injury, as evidenced by the impaired regeneration observed in *Cmg2*^KO^ mice.

### Injury-induced colon regeneration is impaired in *Cmg2*^KO^ mice

To investigate the underlying cellular mechanisms, we performed a detailed analysis of mouse colons after 3 days of DSS withdrawal. In control mice, the crypts were elongated and hypertrophic, a hallmark of the regenerative phase following colitis (Fig. 2a-c) ^25,52,53^. In stark contrast, *Cmg2*^KO^ mice displayed shorter, fewer crypts, and the tissue showed severe alterations. The atrophic crypts in *Cmg2*^KO^ mice had a squamous morphology, differing from the typical columnar shape of epithelial cells seen under normal conditions or during regeneration in *Cmg2*^WT^ mice (Fig. 2a). Additionally, the histological inflammation score was significantly higher in *Cmg2*^KO^ colons during the DSS withdrawal period compared to control mice (Fig. 2d-e).

**Figure 2:**
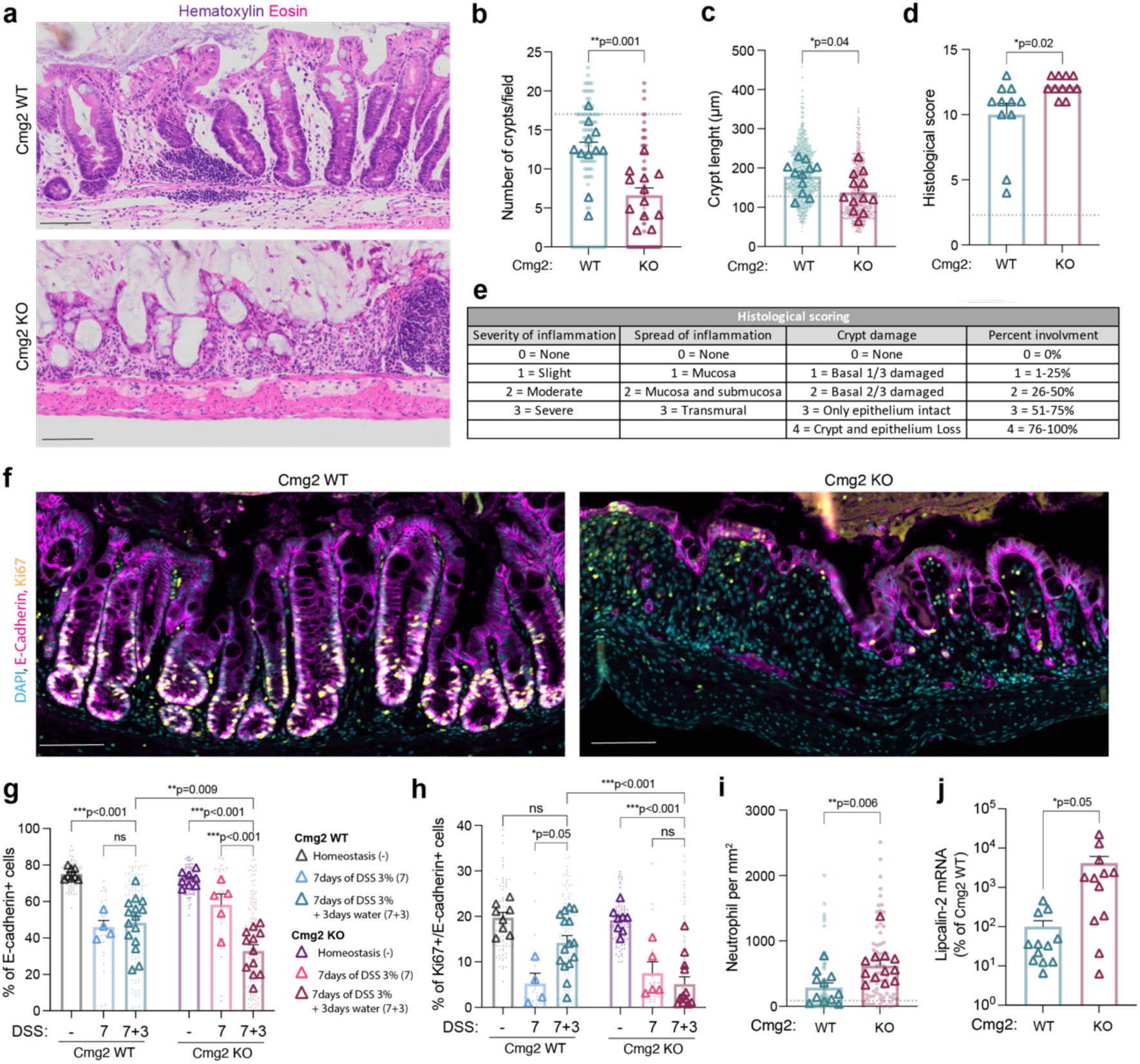
Injury-induced gut regeneration is impaired in *Cmg2*^KO^ mice. 8-week-old *Cmg2*^WT^ or *Cmg2*^KO^ males were given 3% Dextran-Sulfate-Sodium (DSS) in their drinking water for 7 days, then switched to regular drinking water for 3 days to recover. On day10, mice were euthanatized, and their colons were collected. **(a)** Colon tissues were stained for Hematoxylin/Eosin. Representative images of at least n=11 mice per genotype. Scale bar,100µm. **(b)** Number of crypts per field, **(c)** crypt length and **(d-e)** histological score were quantified. Results are mean ± SEM. Each dot represents a single measurement, and each triangle represent the mean per mice. Dotted line represents homeostatic levels of *Cmg2*^WT^ mice. At least n=10 mice per genotype were quantified. *P* values obtained by Two-tailed unpaired t test. **(f)** Colonic sections were stained with anti-ki67, anti-VE-cadherin, and DAPI. Representative images of at least n=10 mice per genotype. Scale bar, 100µm. **(g)** % of E-cadherin+, and **(h)** % of E-cadherin+/Ki67+ cells at homeostasis (-), after 7 days of DSS (7) or 3 days after DSS withdrawal (7+3) were quantified. Results are mean ± SEM. Each dot represents a single measurement and each triangle represent the mean per mice. At least n=4 mice per genotype were quantified. *P* values obtained by Two-way ANOVA Šídák’s multiple comparisons test. **(i)** Immunostaining of neutrophil was performed on colonic section using S100A9 antibody and number of neutrophils per mm2 quantified. Results are mean ± SEM. Each dot represents a single measurement and each triangle represent the mean per mouse. At least n=11 mice per genotype was quantified. *P* values obtained by Two-tailed unpaired t test. In **(j)** qPCR analysis of colonic tissues from *Cmg2*^WT^ and *Cmg2*^KO^ for the intestinal inflammation marker Lipocalin-2. Results are mean ± SEM. n=12 mice per genotype were quantified. *P* values obtained by Two-tailed unpaired t test.

To further examine the damage-induced regeneration process, we assessed epithelial cell proliferation by co-staining for E-cadherin and Ki67 (Fig. 2f). In *Cmg2*^WT^ mice, colons showed elongated crypts with a higher percentage of E-cadherin+/Ki67+ cells compared to the active disease phase (Day 7), returning to levels similar to those seen under control conditions (Day 0) (Fig. 2f, h). In contrast, *Cmg2*^KO^ mice exhibited persistently low levels of proliferating epithelial cells from day 7 to 10, with crypts that remained atrophic. Damage in *Cmg2*^KO^ colons appeared to worsen even after DSS removal, as indicated by a further reduction in epithelial cell numbers (Fig. 2g). Additionally, these colons had more infiltrating neutrophils than those of WT mice (Fig. 2i) and showed increased expression of the inflammation marker lipocalin-2 (Fig. 2j). Collectively, these findings indicate that *Cmg2*^KO^ mice fail to transition from the active disease phase to the regenerative phase after DSS withdrawal, leading to increased damage and impaired repair.

### CMG2 Is Dispensable for YAP/TAZ-Mediated Reprogramming to Fetal-Like Stem Cells

Since *Cmg2*^KO^ mice fail to regenerate their gut upon DSS challenge, we next examined which key steps of the fetal-like regenerative response^25,26,33^ might be affected by Cmg2 depletion. First, we tested whether the expression of Ly6a, a marker of fetal-like stem cells, was influenced by CMG2 in DSS-injured colons. As expected, Ly6a mRNA levels were low under homeostatic conditions and increased significantly 3 days after DSS withdrawal in control mice. (Fig. 3a). Interestingly, this same pattern was also observed in *Cmg2*^KO^ mice (Fig. 3a). Immunofluorescence staining further confirmed the emergence of a fetal-like stem cell population in the colons of both control and *Cmg2*^KO^ mice during regeneration (Fig. 3b-c). Given that this switch to fetal-like stem cells is heavily dependent on YAP/TAZ signaling pathways^25,43^, we investigated whether CMG2 loss affected YAP/TAZ signaling during intestinal regeneration. We measured the mRNA levels of two additional YAP target genes, Cyr61 and CTGF (Fig. 3d-e), both of which were significantly upregulated in the colons of *Cmg2*^KO^ mice after DSS withdrawal (Fig. 3e-f). These findings suggest that CMG2 loss does not impair YAP/TAZ-dependent reprogramming of intestinal cells into fetal-like stem cells.

**Figure 3:**
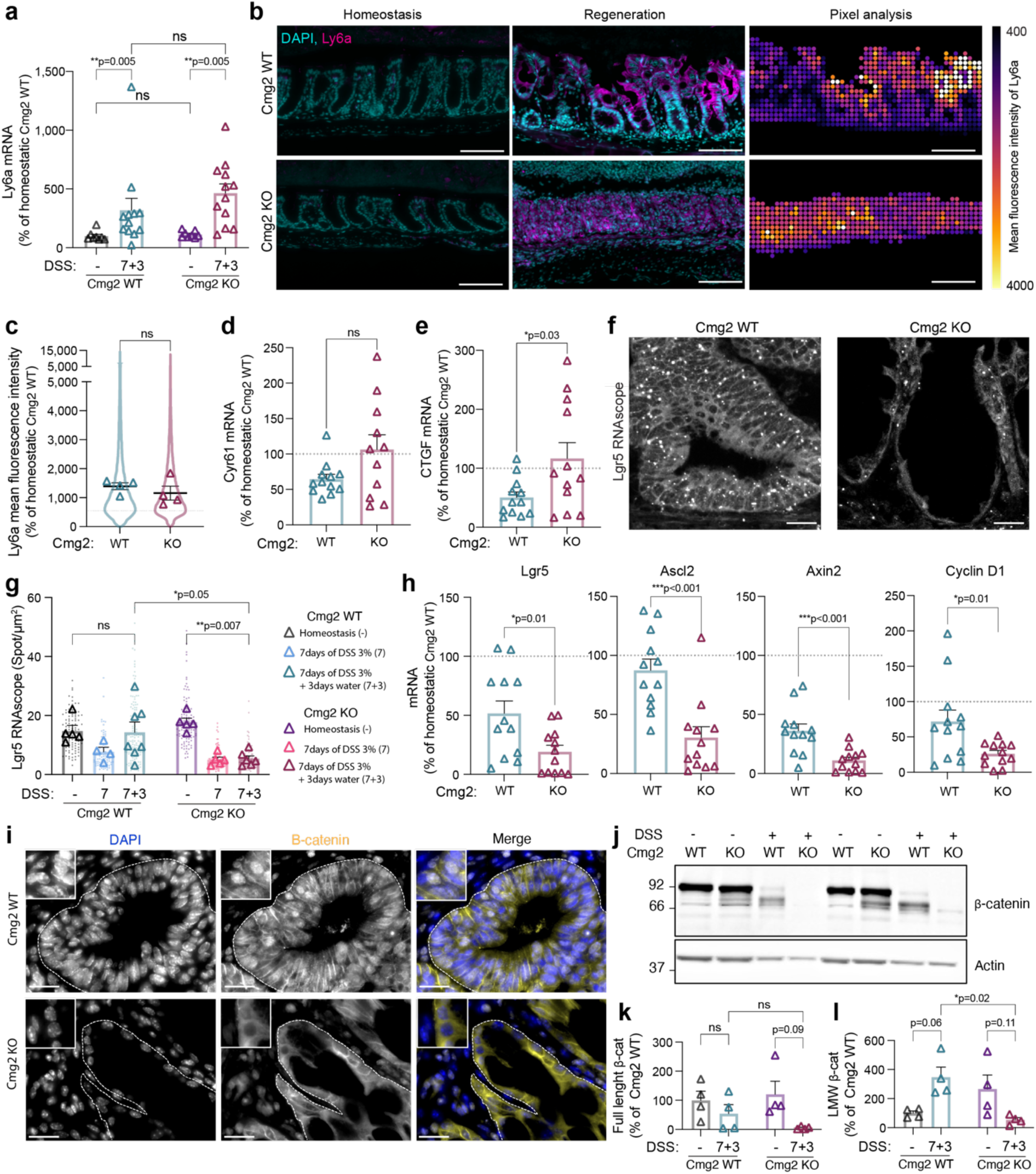
CMG2 is dispensable for YAP/TAZ-dependent reprogramming into fetal-like stem cells but essential for the replenishment of the Lgr5^+^ ISC pool. **(a)** qPCR analysis of colonic tissues from *Cmg2*^WT^ and *Cmg2*^KO^ during homeostasis (-) or 3 days after DSS withdrawal (7+3) for Ly6a. Results are mean ± SEM. Each symbol represents the mean per mouse. At least n=10 mice per genotype were quantified. *P* values obtained by two-way ANOVA Šídák’s multiple comparisons test. **(b-**left and middle panel) Colonic sections were stained with anti-Ly6a, and DAPI and (**b**-right panel) coarse-grain analysis was performed. **(c)** Results are presented as Ly6a mean intensity. Representative images of n=4 mice per genotype. Scale bar, 100µm. Results are mean ± SEM. Dotted line represents homeostatic levels of *Cmg2*^WT^. *P* values obtained by two-tailed unpaired t test. **(d-e)** qPCR analysis of colonic tissues from *Cmg2*^WT^ and *Cmg2*^KO^ 3 days after DSS withdrawal for YAP target genes **(d)** Cyr61 and **(e)** CTGF. Results are mean ± SEM. At least n=11 mice per genotype were quantified. *P* values obtained by two-tailed unpaired t test. **(f-g)** Colonic sections were stained for RNAscope in situ hybridization against Lgr5. Representative images of at least n=4 mice. Scale bar, 100µm. In **(g)** Lgr5 mRNA spots per µm^2^ at homeostasis (-), after 7 days of DSS (7) or 3 days after DSS withdrawal (7+3) were quantified. Results are mean ± SEM. At least n=4 mice per genotype were quantified. *P* values obtained by two-way ANOVA test. **(h)** qPCR analysis of colonic tissues from *Cmg2*^WT^ and *Cmg2*^KO^ 3 days after DSS withdrawal for ISC genes Lgr5, Ascl2, Axin2 and Cyclin D1. Results are mean ± SEM. At least n=12 mice per genotype were quantified. *P* values obtained by two-tailed unpaired t test. **(i)** Colonic sections were stained with anti-β-catenin and DAPI. Representative images of n=4 mice per genotype. Scale bar, 100µm. **(j)** Colon lysates were western blotted against β-catenin and Actin. (**k**) Full-length (95 kDa) or **(l)** low–molecular-weight (LMW) β-catenin/loading control ratio were quantified and normalized to the mean of Cmg2^WT^. Results are mean ± SEM, and n=4 mice per genotype were quantified. *P* values obtained by two-way ANOVA Tukey’s multiple comparisons test.

### CMG2 Is Critical for Restoring the Lgr5+ Intestinal Stem Cell Pool

We then investigated whether the fetal-like stem cells observed in both *Cmg2*^WT^ and *Cmg2*^KO^ mice could successfully transition into adult Lgr5+ ISCs for the replenishment of Lgr5+ stem cells, which is crucial to drive epithelial repair^25,26,33^. After 3 days of DSS withdrawal, Lgr5 expression, monitored by RNAscope, returned to homeostatic levels in the colons of *Cmg2*^WT^ mice, as expected (Fig. 3f-g). In marked contrast, Lgr5 expression remained extremely low in *Cmg2*^KO^ mice, comparable to levels seen during the active disease phase (7 days of DSS treatment) (Fig. 3f-g). qPCR analysis reinforced these findings, showing significantly reduced levels of Lgr5 and three other ISC marker genes (Axin2, Ascl2, and Cyclin D1), all of which are Wnt targets, in *Cmg2*^KO^ mice compared to their wild-type littermates (Fig. 3h). These results indicate that, despite the presence of Ly6a+ fetal-like stem cells, the conversion into Lgr5+ ISCs failed to occur without CMG2.

Following the discovery of the reduced levels of so many Wnt targets, we next investigated the role of CMG2 in modulating Wnt signaling following intestinal damage. Stemness in the intestinal epithelium relies on Wnt signaling^14,54^, which is tightly regulated by various signaling pathways^15,26,44,55–57^, and crucial for effective injury-induced intestinal repair^16^. Initially, we quantified the expression of key molecular components involved in Wnt signaling using qPCR. The mRNA levels of the Frizzled co-receptor LRP6, β-catenin (Fig. S3a-b), and Wnt ligands (Wnt5a, 5b, and 2b) were comparable between the colons of *Cmg2*^WT^ and *Cmg2*^KO^ mice (Fig. S3c), suggesting that CMG2 does not significantly influence the abundance of these major Wnt signaling players.

Next, we analyzed β-catenin activation. Strikingly, while β-catenin was prominently detected in the nuclei of hypertrophic crypts in *Cmg2*^WT^ mice through immunostaining, the nuclei of cells in the atrophic crypts of *Cmg2*^KO^ mice were remarkably devoid of β-catenin (Fig. 3i). Western blot analysis revealed that β-catenin underwent cleavage in the colon of *Cmg2*^WT^ mice 3 days after DSS withdrawal (Fig. 3j-l), a hallmark of enhanced transcriptional activation^58^. In contrast, *Cmg2*^KO^ mice exhibited extremely low levels of both full-length and cleaved β-catenin during DSS treatment (Fig. 3j-l), despite displaying normal levels under basal conditions (Fig. S3d-e). Collectively, these findings demonstrate that CMG2 is essential for the reactivation of Wnt signaling, facilitating the transition from fetal-like to adult ISCs necessary for effective intestinal repair.

## DISCUSSION

The present study underscores a critical and previously unknown role of CMG2 in gut function. The most severe form of HFS is often fatal due to intractable diarrhea. Our results show that *Cmg2*^KO^ mice exhibited no abnormalities in the colon under normal conditions, correlating with normal function of enterocytes derived from HFS patient cells^13^. Differences became apparent when the mice were subjected to chemical induced colitis. While both control and *Cmg2*^KO^ mice showed similar disease severity, the knockout mice failed to regenerate their colons after DSS withdrawal, resulting in persistent bloody diarrhea and significant weight loss—symptoms that closely mirror those seen in HFS patients.

Our work demonstrates the implication of CMG2 in intestinal regeneration after damage, a process that is thought to occur in at least two phases. CMG2 was found to be non-essential during the initial de-differentiation phase, characterized by high YAP activation, with repression of Wnt signaling and ISC gene expression^44,55,56^, since we observed that fetal-like stem cells were present in the damaged colon of both *Cmg2*^WT^ and *Cmg2*^KO^ mice. Our data show that CMG2 is crucial for the second phase of Wnt- and Ascl2-dependent reacquisition of Lgr5+ stemness^59^, thereby enabling the restoration of intestinal homeostasis after injury. The fact that CMG2 is not required for basal gut homeostasis but is essential for replenishing the ISC pool aligns with the idea that Lgr5+ cells are dispensable for maintaining normal epithelial function^24,60^. During intestinal regeneration, the transition of these fetal-like stem cells to adult ISCs, require the downregulation of YAP and reactivation of Wnt signaling^61–66^, suggesting a major role of Cmg2 in tunning these signaling pathways.

Our findings expand the growing network of connections between CMG2 and Wnt signaling pathways, as well as the connection to stem cells. Previous work on CMG2 identified it as an interactor of Wnt signaling co-receptor LRP6 through studies on anthrax toxin receptors^62^. We confirmed this interaction, demonstrating that LRP6 and CMG2 co-internalize during toxin entry^18^. CMG2 have also been shown to modulate LRP6 levels in a concentration-dependent manner, where both low and high levels of CMG2 reduce LRP6 abundance in cultured cells^18^. In mouse 3T3-L1 cells, downregulation of CMG2 consistently impaired Wnt-induced β-catenin stabilization^18^. CMG2 not only connects to canonical LPR6-dependent Wnt signaling. We indeed found that during zebrafish embryonic development, CMG2 is necessary for the non-canonical Wnt-dependent oriented cell division of epiblast cells, necessary for embryo extension^17,21^, a process that still involved LRP6. Recent studies in cancer biology extended the connection of CMG2-Wnt to stem cells^20^. CMG2 levels correlated with increased expression of stemness-related genes, such as Lgr5, as well as enhanced self-renewal and metastatic potential^20^. Manipulating CMG2 levels through over-expression or silencing altered nuclear β-catenin, consistent with previous observations in 3T3-L1 cells and our current findings that Cmg2 deficiency impaired β-catenin activation and Lgr5+ ISC in the damaged. Noteworthily, the absence of CMG2 does not result in the same effects as LRP6 or Wnt deficiencies, which can cause early lethality or milder phenotypes^67^. This indicates that CMG2 modulates Wnt signaling in a context-specific manner within the gut.

In conclusion, our study demonstrates that CMG2 is a key player in gut regeneration, specifically impacting the Wnt-mediated replenishment of the Lgr5+ stem cell pool following injury. Combined with its established role in gastric stem cells, this work suggests a broader function for CMG2 in stem cell biology, particularly in tissue regeneration involving fetal-like reversion, as also seen in the stomach^66,68,69^. From a clinical perspective, our findings provide a molecular basis for the persistent diarrhea seen in HFS patients. In HFS infants, a diarrheal event may lead to failure in replenishing the stem cell pool, thus preventing reestablishment of epithelial barrier. These insights suggest that HFS patients might benefit from therapeutic approaches currently used for chronic diarrhea conditions, such as inflammatory bowel diseases (IBD).

## MATERIAL AND METHODS

### Animals

CMG2 knockout mice were generated by targeted deletion of the gene exon 3 as described previously^2^. Cmg2 mutant mice were generated and backcrossed for at least 10 generations onto the C57Bl6/J genetic background (Charles River Laboratories). *Cmg2*^HET^ mice were kept in a specific pathogen free (SPF) environment with 12 h light and 12 h dark cycle with ad libitum access to food and water. Experiments were performed using littermate from the different genotypes and animals housed depending on their genotype. During the conduct of DSS experiment and during all data analysis, mice genotype was blinded. Characteristics of animals used in this study as well as animal care and monitoring during DSS experiment are described in Supplementary information. For animal experimentation, all procedures were performed according to protocols approved by the Veterinary Authorities of the Canton Vaud and according to the Swiss Law (License VD 3497, EPFL) and were in accordance with the ARRIVE guidelines and 3R principle (Reduction, Replacement, Refinement) for laboratory animals.

### DSS Induced colitis

Eight-weeks *Cmg2*^WT^ or *Cmg2*^KO^ males were given 3% Dextran Sulfate Sodium (MP Biomedicals:160110) in the drinking water for 7 days, then switched to regular drinking water for 3 days and allowed to recover. Control Eight-weeks *Cmg2*^WT^ or *Cmg2*^KO^ males were given drinking water that did not contain DSS. Water consumption was similar in DSS-treated or control mice. During DSS treatment and recovery, mice were weighed, scored daily and euthanized if they developed severe disease based on the scoring criteria described in Supplementary information (weight loss, coat appearance, eyes/nose discharge, breathing, activity and posture). Mice were housed in individual cage for 30-60min daily in order to collect mice feces. Feces were either stored at -20° for further analyses or diluted at 100mg/ml in PBS 0,1% Tween-20 and used to evaluate the presence of occult blood using urine test strip (Multistix 8SG). Daily evaluation of Disease activity Index (DAI) (described in Supplementary information) was performed and based on stool consistency, presence of occult blood and body weight loss. On day 10, mice were euthanized with injection of pentobarbital. Colon was collected, measured and weighted. Tissues were divided in two, the first part being store at -80° for RNA and protein extraction and the second part fixed in 4% paraformaldehyde for histological studies.

### Histological studies

Tissues collected after euthanasia were fixed in 4% paraformaldehyde overnight at 4°, embedded in paraffin and sectioned at 4µm thickness. Sections were deparaffinized with xylene and rehydrated with distilled water through a series of graded alcohol. Tissues were stained with hematoxylin & eosin (H&E), Sirius Red or Alcian blue, using standard protocols. Each colon was scored by a pathologist who assigned four scores based on severity and spread of inflammation, crypt damage and percent involvement. For immunohistochemistry, de-waxed samples were subjected to antigen retrieval by boiling in citrate buffer (10 mM, pH 6.0) for 20 min, except for Sca1/Ly6a immunostaining which was subjected to antigen retrieval by treating Proteinase K (20ug/ml) for 10min at room temperature. Non-specific antigenic sites were blocked with 1% bovine serum albumin (BSA) and samples were hybridized with primary antibody (Listed in the antibody section) in a humidity chamber at 4 °C overnight. After washing, samples were incubated with secondary antibodies (Listed in antibody section) for 1h at room temperature. After washing, tissues were counterstained with DAPI (1/5000 dilution) and mounted with Prolong Gold Antifade Mounting medium (Thermo Fisher, P36930).

### Antibodies

Actin (Millipore: MAB1501: RRID: AB_2223041; mouse: 1:4000 dilution).

Collagen-VI (Abcam: ab6588: RRID:AB_305585; rabbit: 1:1000 dilution)

E-cadherin (Abcam: ab76055: RRID:AB_1310159; mouse: 1:500 dilution)

Ki-67 (Abcam: ab16667: RRID:AB_302459; rabbit: 1:1000 dilution)

S100A9 (neutrophil) (Novus Biological: NB110-89726: rabbit: 1:200 dilution)

B-catenin (For IF: Sigma: C2206: RRID:AB_476831: Rabbit: 1/200 dilution)

B-catenin (For WB: BD biosciences: 610154: RRID:AB_397555: Rabbit: 1/1000 dilution)

Sca1/Ly6a (Abcam: ab51317: RRID_AB1640946: Rat: 1/200 dilution)

Mouse-HRP (GE Healthcare: NA931V; RRID: AB_772210; mouse: 1: 3000 dilution).

Rabbit-HRP (GE Healthcare: NA934V; RRID: AB_772206; rabbit: 1: 3000 dilution).

Mouse-Alexa488 (ThermoFisher Scientific: A-11029; RRID: AB_2534088; 1:1000 dilution).

Mouse-Alexa568 (ThermoFisher Scientific: A-11037; RRID: AB_2534013; 1:1000 dilution).

Mouse-Alexa647 (ThermoFisher Scientific: A-31571; RRID: AB_162542; 1:1000 dilution).

Rabbit-Alexa488 (ThermoFisher Scientific: A-21206; RRID: AB_2535792; 1:1000 dilution).

Rabbit-Alexa568 (ThermoFisher Scientific: A-11042; RRID: AB_2534017; 1:1000 dilution).

Rabbit-Alexa647 (ThermoFisher Scientific: A-31573; RRID: AB_253183; 1:800 dilution).

Rat-Alexa568 (ThermoFisher Scientific: A-11077; RRID: AB_2534121; 1:1000 dilution).

DAPI (ThermoFisher Scientific: D1306; RRID: AB_2629482; 1:5000 dilution).

Hoechst (Sigma: 94403; 1:5000 dilution).

### RNAscope

RNAscope Multiplex Fluorescent V2 assay (Bio-techne, Cat. No. 323110) was performed according to manufacturer’s protocol on 4 um paraffin sections with 15 minutes target retrieval at 95°C and Protease III 30 minutes at 40°C, hybridized with the probes Mm-Lgr5-C3 (Bio-techne, Cat. No. 312171-C3), Mm-Ppib-C1 (Bio-techne, Cat. No. 313911) as positive control and DapB-C1 (Bio-techne, Cat. No. 310043) as negative control at 40°C for 2 hours and revealed with TSA Opal570 (Akoya Biosciences, Cat. No. FP1488001KT). Tissues were counterstained with DAPI and mounted with Prolong Gold Antifade Mounting medium (Thermo Fisher, P36930).

### Image acquisition and analyze

Brightfield and fluorescent images were acquired on an Olympus VS200 whole slide scanner using OlyVIA software and 20x air objective or 40x air objective. Detailed information about technical specifications in Supplementary information.

For all analysis, annotations areas were selecting manually using DAPI channel exclusively (Supplementary Fig. 3a-b). During all analysis, mice genotype and channel of interest were blinded. Analysis and quantification of images was performed using the QuPath software^70^. Analysis workflows are available in Supplementary figure 3. Briefly, the percentage of Collagen VI area per tissue section was quantified using pixel classifier/thresholder for the Col6 channel (Supplementary Fig. 3c). The number of Lgr5 RNAscope spots per area was quantify using the RNAscope code detailed in Supplementary Fig. 3d). The percentage of positive cells positive to different marker (E-cadherin, Ki67 or S100A9), was quantified using the STARDIST Qupath extension for cell detection (https://github.com/stardist/stardist) and specific classifiers were created for each marker (Supplementary Fig. 3e). To determine Ly6a intensity, a coarse grain analysis of Ly6a immunostaining was performed (Supplementary Fig. 3f).

### Quantification of fecal and serum lipocalin-2 by ELISA

Frozen fecal samples collected at day 0, 4, 7 and 9 during DSS induced colitis experiment, were reconstituted in PBS containing 0.1% Tween-20 (100mg/ml), vortexed at 4° for 20min and centrifuged 10min at 12,000rpm at 4°. Clear supernatants were collected and stored at -20° until analysis. Fecal lipocaline-2 was quantified using the Duoset murine Lcn-2 ELISA kit (R&D Systems) accordingly to manufacture instructions.

### qRT-PCR

Mice tissues were disrupted using Lysing Matrix tube (MP Biomedicals) and homogenized in a tissue lyser (Qiagen) 2 x 2min, followed by RNA extraction using RNA easy mini extraction kit (Qiagen). RNA concentration was measured by spectrometry and 1mg of total RNA was used for reverse transcription. We then used a 1/10 dilution of the cDNA to perform quantitative real-time PCR using SYBR MasterMix (Life Technology) on a Quantstudio 6 Real-Time PCR System (Applied Biosystem). mRNA level in triplicate was normalized using Ribosomal protein S9 (RSP9) and Eukaryotic Translation Elongation Factor 1 Alpha 1 (EEF1A1) and Results were expressed as 2^(-DDCt)*100%.

Primers (all 5’ to 3’) used are listed below:

Mouse *Col6a1*: F: TGCCCTGTGGATCTATTCTTCG; R: CTGTCTCTCAGGTTGTCAATG

Mouse *Axin2*: FOR: TGACTCTCCTTCCAGATCCCA; R: TGCCCACACTAGGCTGACA

Mouse *Lgr5*: F: AGGGTGGACTGCTCCGACCTG; R: AAGACGTAACTCCTCCAGGAAGCGG

Mouse *CyclinD1*: F: TCAAGTGTGACCCGGACTG; R: ACTCCAGAAGGGCTTCAATCT

Mouse *Ascl2*: F: TTTCCTGTGCCGCACCAGAACT; R: CAGCGACTCCAGACGAGGTGG

Mouse *Tnfα*: F: TCCACTTGGTGGTTTGCTACG; R: ATGAGCACAGAAAGCATGATC

Mouse *Ifnγ*: F: GCCACGGCACAGTCATTGA; R: TGCTGATGGCCTGATTGTCTT

Mouse *IL10*: F: CTTACTGACTGGCATGAGGATCA; R: GCAGCTCTAGGAGCATGTGG

Mouse *IL1β*: F: CAACCAACAAGTGATATTCTCCATG; R: GATCCACACTCTCCAGCTGCA

Mouse *IL6*: F: CTGCAAGAGACTTCCATCCAG; R: AGTGGTATAGACAGGTCTGTTGG

Mouse *IL4*: F: GGTCTCAACCCCCAGCTAGT; R: GCCGATGATCTCTCTCAAGTGAT

Mouse *Tgfβ*: F: CACCGGAGAGCCCTGGATA; R: TGTACAGCTGCCGCACACA

Mouse *Lipocalin-*2: F: AAGGCAGCTTTACGATGTACAGC R: CTTGCACATTGTAGCTGTGTACC

Mouse *Ly6a*: F: CCTACCCTGATGGAGTCTGTGT; R: CACGTTGACCTTAGTACCCAGG

Mouse *Ctgf*: F: CTGCGCTAAACAACTCAACGA; R: GCAGATCCCTTTCAGAGCGG

Mouse *Lrp6*: F: GGATGCCCTGCCCACTACTC; R: CAGAAGAGCGCCATCAACCGC

Mouse β*-catenin*: F: CTGGCCGTATCCACCAGAGT; R: GAAGCGGCTTTCAGTCGAGC

Mouse *Wnt5a*: F: TGTCTTCGCACCTTCTCCAATG; R: CTCCTTCGCCCAGGTTGTTATAG

Mouse *Wnt5b*: F: TCCTGGTGGTCACTAGCTCTG; R: TGCTCCTGATACAACTGACACA

Mouse *Wnt2b*: F: CGTTCGTCTATGCTATCTCGTCAG; R: ACACCGTAATGGATGTTGTCACTAC

Mouse *Rsp9*: F: GACCAGGAGCTAAAGTTGATTGGA; R: TCTTGGCCAGGGTAAACTTGA

Mouse *Cox6a1*: F: CTCTTCCACAACCCTCATGTGA; R: GAGGCCAGGTTCTCTTTACTCATC

Mouse *Eef1a1*: F: TCCACTTGGTCGCTTTGCT; R: CTTCTTGTCCACAGCTTTGATGA

### Western blot

Mice tissues were lysed 30 min at 4°C on rotating wheel in IP buffer (0.5% NP40; 500mMtris-HCl, pH=7.4; 20mM EDTA; 2mM benzamidine; 10mM NaF and a cocktail of protease inhibitors). The suspended tissues were then put in a Lysing Matrix tube (MP Biomedicals) and homogenized in a tissue lyser (Qiagen) 2 x 2min at 20Hz. Homogenates were then pelleted at 5,000 r.p.m. for 3 min at 4 °C. Supernatants were subjected to preclearing with G sepharose beads and protein amount was quantified by BCA assay (Pierce). After protein concentration normalization, samples were boiled in Laemmli buffer for 5 min before analysis by SDS–PAGE using 4–20% Bis-Tris gradient gels under reducing condition and western blotting with rabbit anti-collagenVI antibody and mouse anti-Actin antibody.

### Quantification and statistical analysis

Unless otherwise stated, each data point corresponds to a single measurement and each triangle correspond to the mean per mice. Statistical analysis was carried out using Prism software. Data representations and statistical details can be found in the description of the Figures. For ANOVA analysis, *P* values were obtained by post hoc tests to compare every mean and pair of means (Tukey’s & Sidak’s).

### Data and Code availability

All details regarding codes used for immunofluorescence analysis are available in GitHub (https://github.com/upvdg) and will be deposited at Zenodo and publicly available as of the date of publication.

## ACKNOWLEDGEMENTS

We would like to thank the EPFL Facilities (BioImaging & Optics Core Facility, Center of PhenoGenomics and Histology Core Facility) and all the members of the F.G.v.d.G. lab for discussions and suggestions. This work was supported by the Swiss National Science Foundation, by the Gelù Foundation and by the Foundation Les Mûrons.

## AUTHOR CONTRIBUTIONS

LB, AC, and BK, conducted the experiments, acquired and analyzed the data. OB and RG developed codes used for immunofluorescence analysis. LB and GG conceptualized the study and wrote the manuscript. AC and BK critically reviewed the manuscript.

## DECLARATION OF GENERATIVE AI AND AI-ASSISTED TECHNOLOGIES IN THE WRITING PROCESS

During the preparation of this work the author(s) used ChatGPT in order to improve the English of the text. After using this tool/service, the author(s) reviewed and edited the content as needed and take(s) full responsibility for the content of the publication.

---

**Figure S1:**
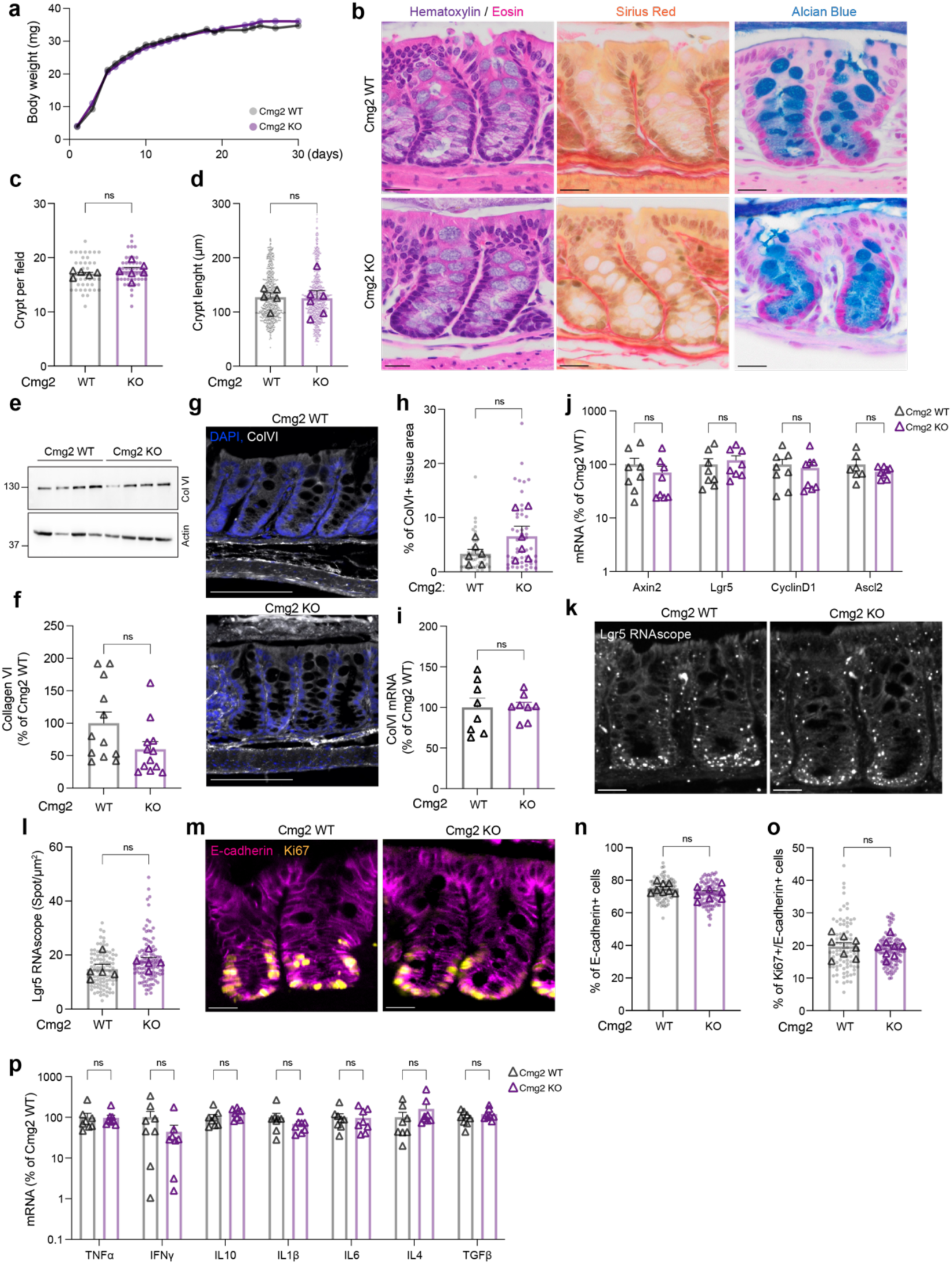
*Cmg2* KO mice have normal guts under basal conditions. **(a)** Body weight of Cmg2^WT^ and Cmg2^KO^ over a 30weeks period. Results are mean ± SEM. Each dot represents the mean of at least n=8 mice per genotype. (**b**) Colon tissues under basal conditions, from 8-weeks-old *Cmg2*^WT^ and *Cmg2*^KO^ mice were stained for Hematoxylin/Eosin, Sirius Red and Alcian Blue. Representative images of at least n=8 mice per genotype. Scale bar, 20µm. (**c**) Number of crypts per field, (**d**) crypt length were quantified and showed as superplot. Results are mean ± SEM. Each dot represents a single measurement and each triangle represent the mean per mouse. At least n=5 mice per genotype were quantified. *P* values obtained by unpaired two-tailed *t* test. (**e**) Colon lysates were analyzed by SDS–PAGE using 4–12% Bis-Tris gradient gels under reducing condition and western blotted against all the collagen VI. Migration of the molecular weight markers (in kDa) are indicated on the left. (**f**) CollagenVI/loading control ratio were quantified and normalized to the mean of Cmg2^WT^. Results are mean ± SEM, and n=12 mice per genotype were quantified. *P* values obtained by unpaired two-tailed *t* test. (**g**) Colon tissues were immunostained with anti-collagenVI and DAPI. Representative image of n=6 mice per genotype. Scale bar, 20µm. **(h)** % of Collagen VI area per tissue section was quantified on n=6 mice per genotype. Results are mean ± SEM. Each dot represents a single measurement and each triangle represent the mean per mouse. *P* values obtained by unpaired two-tailed *t* test. q PCR analysis of colonic tissues for (**i**) Col6a1, and (**j**) ISC markers Lgr5, Ascl2, Axin2 and CyclinD1 genes were performed on n=8 mice per genotype. *P* values obtained by unpaired two-tailed *t* test in (i) and by two-way ANOVA Šídák’s multiple comparisons test in (j). (**k**) Colonic tissues were stained for RNAscope in situ hybridization against Lgr5 and **(l)** number of Lgr5 mRNA spots per µm^2^ were quantified. (**m**) Colonic sections were stained anti-ki67, anti-E-cadherin and DAPI. (**n**) % of E-cadherin+ epithelial cells, and (**o)** % of E-cadherin+/Ki67+ epithelial cells, were quantified on n=8 mice per genotype. *P* values obtained by unpaired two-tailed *t* test. (**p**) qPCR analysis of colonic tissues for inflammatory cytokines genes genes were performed on n=8 mice per genotype. *P* values obtained by two-way ANOVA Šídák’s multiple comparisons test.

**Figure S2:**
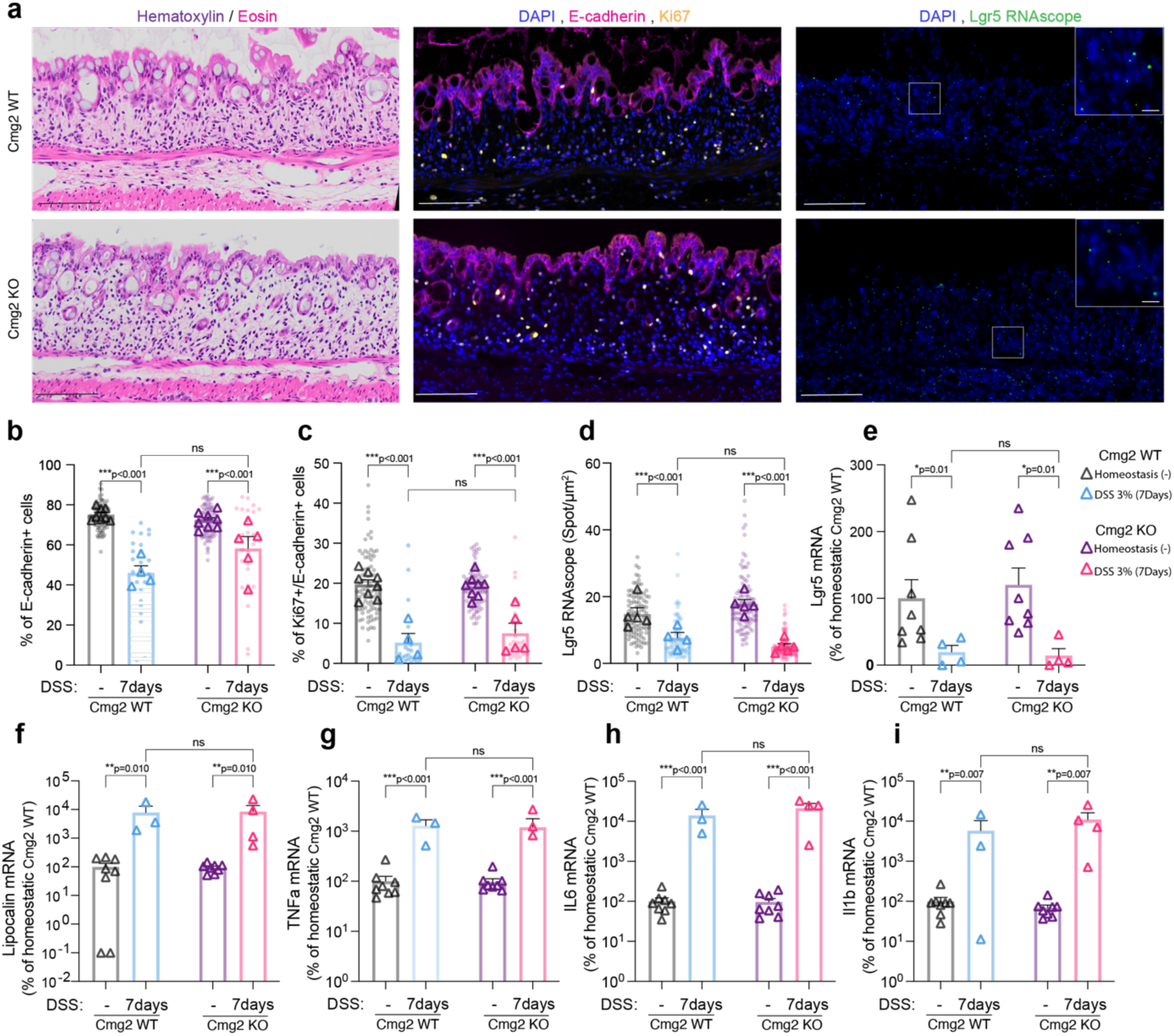
CMG2 loss does not enhance the severity of DSS-induced colitis. 8-week-old *Cmg2*^WT^ or *Cmg2*^KO^ males were given 3% Dextran-Sulfate-Sodium (DSS) in their drinking water for 7 days. On day7, mice were euthanatized and colon collected. (**a**) Colon tissues were stained for Hematoxylin/Eosin (left panel), stained with anti-ki67, anti-VE-cadherin and DAPI (middle panel) or stained for RNAscope in situ hybridization against Lgr5 (right panel). Representative images of at least n=4 mice per genotype. Scale bar,100µm for main image and 10um for magnification. **(b)** % of E-cadherin+ cells, **(c)** % of E-cadherin+/Ki67+ cells, and **(d)** number of Lgr5 mRNA spots per µm^2^, were quantified. Results are mean ± SEM. Each dot represents a single measurement and each triangle represent the mean per mouse. At least n=4 mice per genotype were quantified. *P* values obtained by two-way ANOVA Šídák’s multiple comparisons test. **(e-i)** Quantitative real-time PCR analysis of colonic tissues from *Cmg2*^WT^ and *Cmg2*^KO^ for **(e)** Lgr5, **(f)** Lipocalin-2 **and (g-i)** inflammatory cytokines genes. Results are mean ± SEM. Each symbol represents a single measurement. At least n=3 mice per genotype were quantified. *P* values obtained by two-way ANOVA Šídák’s multiple comparisons test.

**Figure S3:**
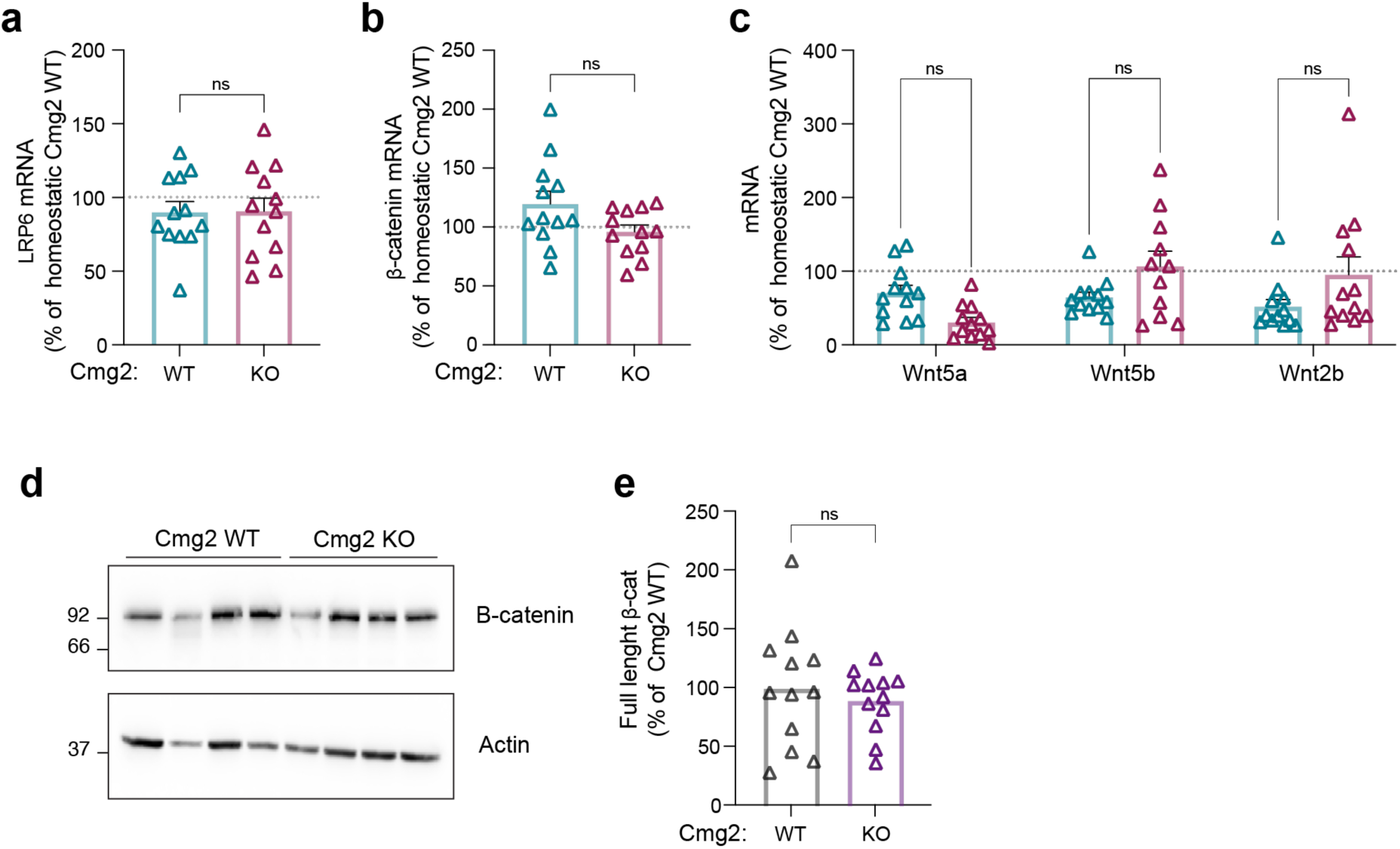
CMG2 does not affect Wnt ligand secretion or expression of LRP6 and B-catenin. **(a-c)** qPCR analysis of colonic tissues from Cmg2^WT^ and Cmg2^KO^ mice 3 days after DSS withdrawal (day10) for **(a)** LRP6, **(b)** β-catenin and **(c)** Wnt ligand, Results are mean ± SEM. Each symbol represents the mean per mouse. dotted line represents the mean of homeostatic Cmg2^WT^ mice. n=12 mice per genotype was quantified. *P* values obtained by two-tailed unpaired t test in (a-b) or by Two-way ANOVA Šídák’s multiple comparisons test in (c). **(d)** Colon lysates were western blotted against β-catenin. **(e)** β-catenin/loading control ratio were quantified and normalized to the mean of Cmg2^WT^. Results are mean ± SEM, and n=12 mice per genotype were quantified. Each triangle represents the mean per mouse. *P* values obtained by unpaired two-tailed *t* test.

**Figure S4:**
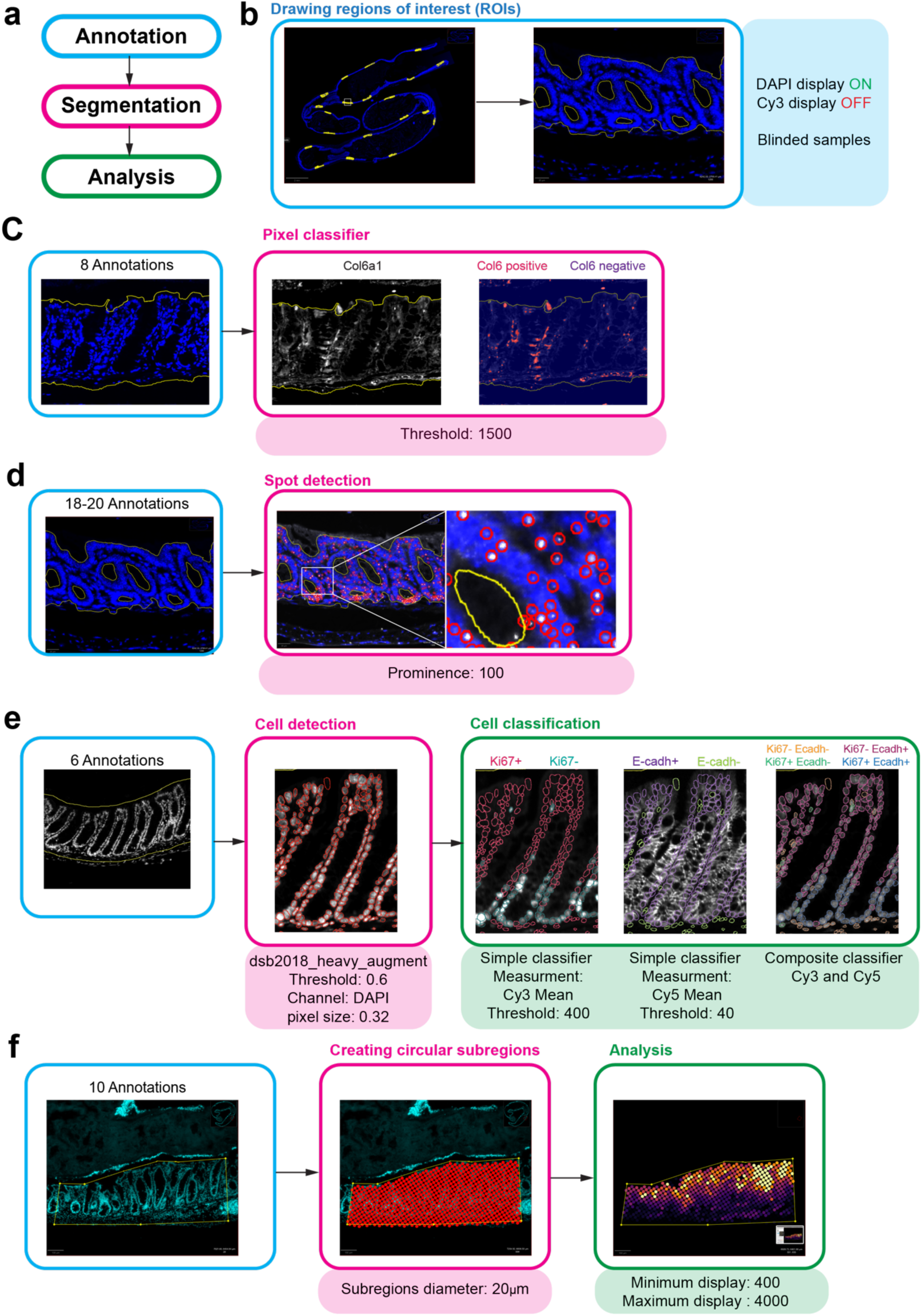
Microscopy workflow: **(a)** Microscopy workflow follow three different steps including annotation, segmentation and analysis. **(b)** Annotation of region of interest is performed on blinded samples (blind genotype and mice id). During this step, channels of interest are turned off and annotations are drawn manually using the DAPI channels. 6 to 20 annotations per sample are drawn along the whole tissue section. **(c)** To quantify the percentage of Col6a1+ tissue area, annotations are drawn as described and Qupath pixel classifier is performed. Threshold is set by visual inspection and applied to all samples. **(d)** For RNAscope spot detection, annotations are drawn and script RNAscope_Count has been used to quantify the number of spots per annotation area. Prominence is set by visual inspection and apply to all samples. **(e)** To quantify the percentage of E-Cadherin+ and Ki67+ cells, annotations were drawn and cells detected using DAPI channel and Qupath STARDIST. Pixel classifiers were then used to classify cells accordingly to E-cadherin and Ki67 staining. **(f)** To analyze Ly6a in colon section, annotations were divided in multiple circular subregions of 20um diameter. The mean fluorescence intensity of Ly6a in each subregion was then quantify and shown using Qupath display.

## Supplementary information

### Animal’s characteristics

Animals involve in DSS experiments:

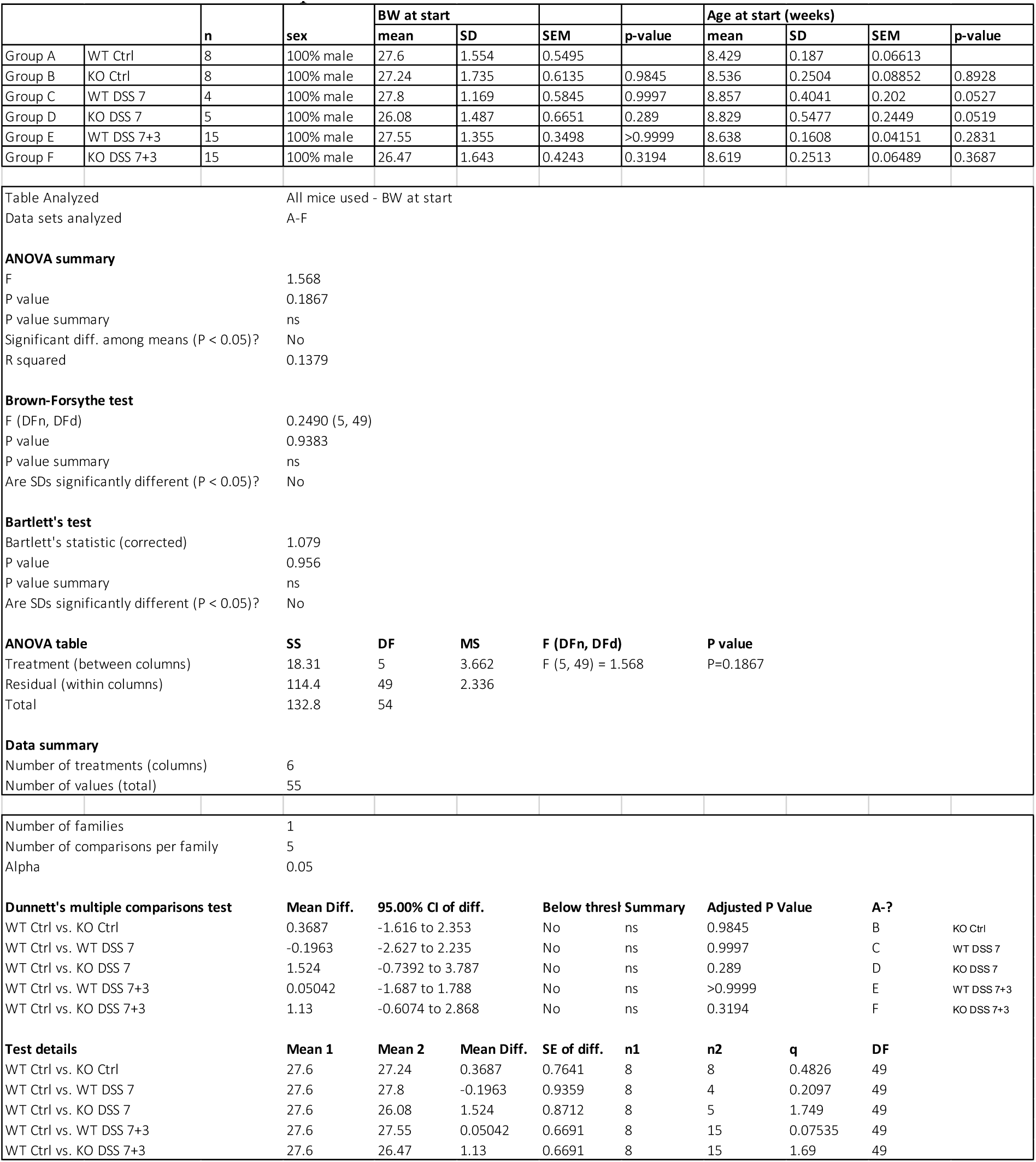

Other mice used in this study:

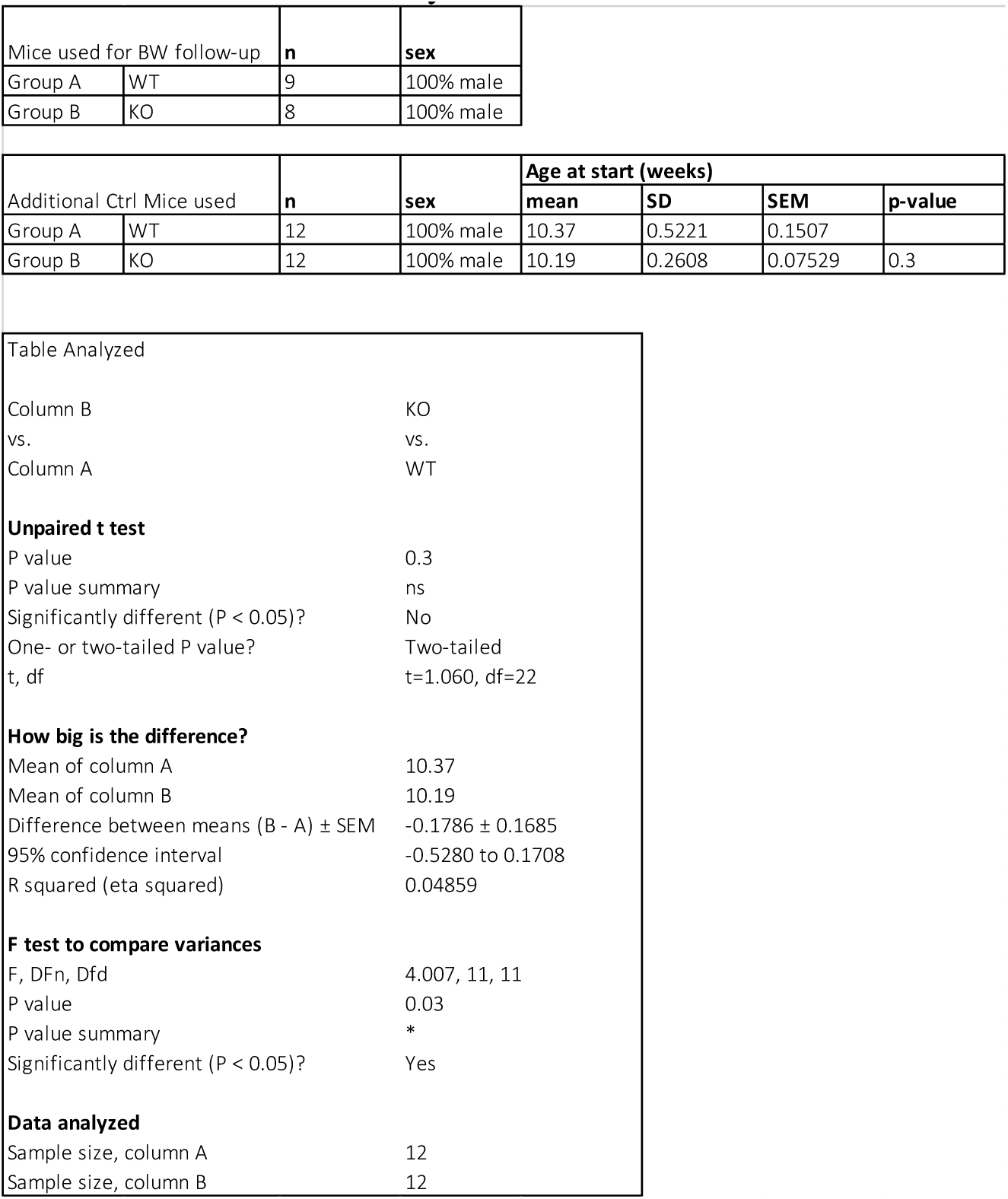

### Animal care and monitoring

During DSS experiment, animals care and welfare was monitored accordingly to the following scoresheet validated by the approved by the Veterinary Authorities of the Canton Vaud and according to the Swiss Law (License VD 3497, EPFL).

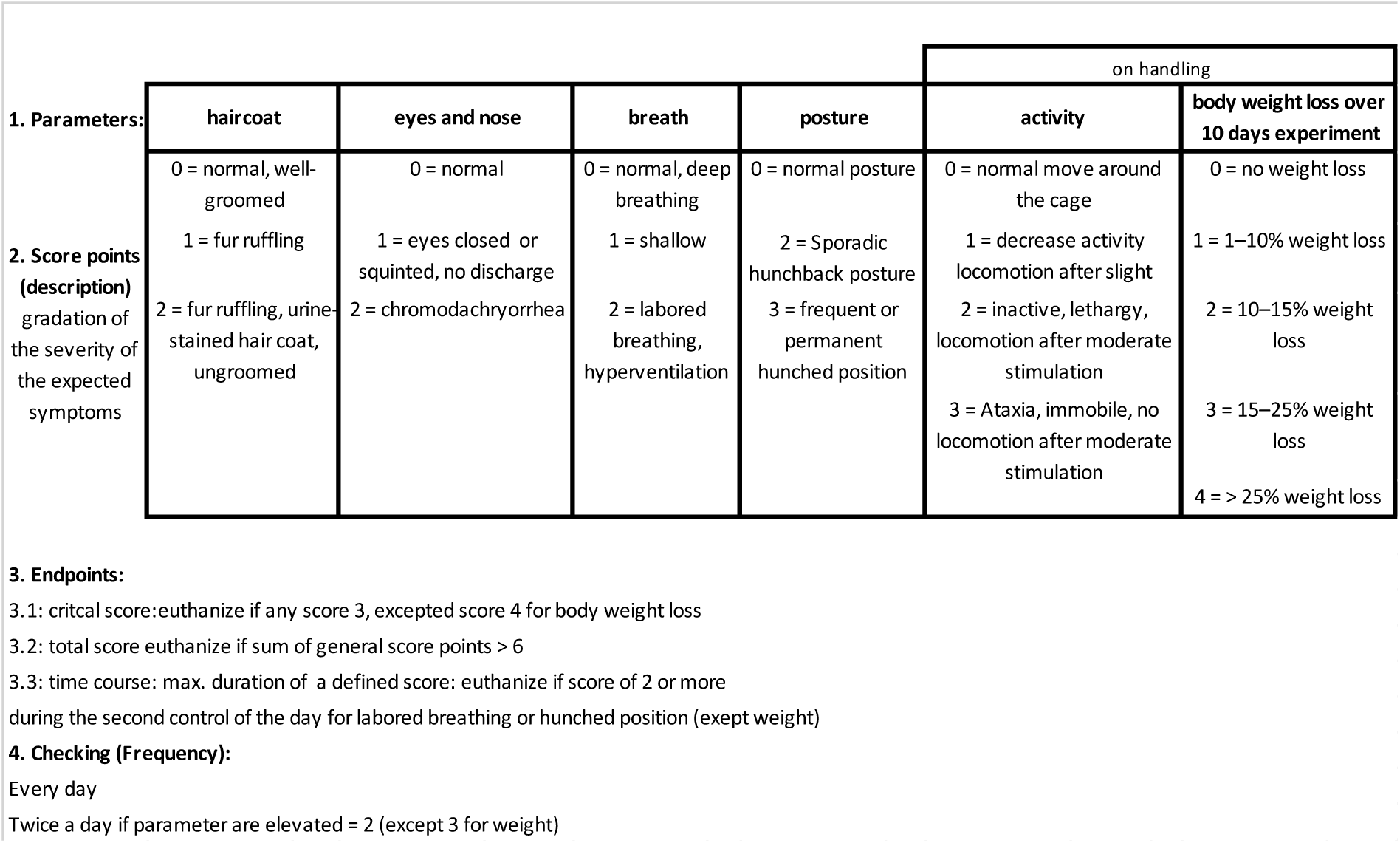

### Microscope technical specifications

|  |  |  |
| --- | --- | --- |
| Cameras | <u>Color UI 3280 CP</u> | <u>Black/White VS20-304M</u> |
| Chip Technology | CMOS Color | CMOS Grayscale |
| Sensor Format | 2456 x 2054 | 4104 x 3006 |
| Chip size | 8.47mm x 7.08mm | 14.15mm x 10.37mm |
| Pixel size | 3.45 um x 3.45 um | 3.45 um x 3.45 um<br>(used at 2x binning) |
| Dynamic Range | 8-bit | 12-bit |
| C-mount | 0.63x | 1.0x |

Objectives
| Objective | Mag/NA | Medium | Contrast | WD (mm) |
| --- | --- | --- | --- | --- |
| PLAPON | 2x/0.08 | air | Pol | 6.20 |
| UPLXAPO | 4x/0.16 | air | Pol | 13.00 |
| UPLXAPO | 10x/0.40 | air | Pol | 3.10 |
| UPLXAPO | 20x/0.80 | air | Pol | 0.60 |
| UPLXAPO | 40x/0.95 | air | Pol | 0.18 |
| UPLXAPO | 40x/1.40 | oil | Pol | 0.13 |

| Name | Excitation | Dichroic | Emission | Nº |
| --- | --- | --- | --- | --- |
| <u>DAPI</u> | BP 378/52 | Pentaband BP | BP 432/36 | <u>FF01-378/52-25</u><br><u>FF01-432/36-25</u> |
| <u>FITC</u> | BP 474/27 | Pentaband BP | HC 515/30 | <u>FF01-474/27-25</u><br><u>FF01-515/30-25</u> |
| <u>Cy3</u> | BP 554/23 | Pentaband BP | HC 595/31 | <u>FF01-554/23-25</u><br><u>FF01-595-31-25</u> |
| <u>Cy5</u> | BP 635/18 | Pentaband BP | BP 685/40 | <u>FF01-635/18-25</u><br><u>FF02-685/40-25</u> |
| <u>Cy7</u> | BP 735/28 | Pentaband BP | BP 809/81 | <u>FF01-735/28-25</u><br><u>FF02-809/81-25</u> |

Full spectrum for all filters

